# Germline genetic variants associated with leukocyte-genes predict tumor recurrence in breast cancer patients

**DOI:** 10.1101/312355

**Authors:** Jean-Sébastien Milanese, Chabane Tibiche, Jinfeng Zou, Pengyong Han, Zhi Gang Meng, Andre Nantel, Simon Drouin, Richard Marcotte, Edwin Wang

**Author notes:** Corresponding to: E.W., Department of Biochemistry & Molecular Biology, Medical Genetics, and Oncology, University of Calgary, 3330 Hospital Dr. NW, Calgary, Canada, T2N 4N1.

## Abstract

Germline genetic variants such as BRCA1/2 play an important role in tumorigenesis and clinical outcomes of cancer patients. However, only a small fraction (i.e., 5-10%) of inherited variants has been associated with clinical outcomes (e.g., BRCA1/2, APC, TP53, PTEN and so on). The challenge remains in using these inherited germline variants to predict clinical outcomes of cancer patient population. In an attempt to solve this issue, we applied our recently developed algorithm, eTumorMetastasis, which constructs predictive models, on exome sequencing data to ER+ breast (n=755) cancer patients. Gene signatures derived from the genes containing functionally germline genetic variants significantly distinguished recurred and non-recurred patients in two ER+ breast cancer independent cohorts (n=200 and 295, P=1.4×10^−3^). Furthermore, we found that recurred patients possessed a higher rate of germline genetic variants. In addition, the inherited germline variants from these gene signatures were predominately enriched in T cell function, antigen presentation and cytokine interactions, likely impairing the adaptive and innate immune response thus favoring a pro-tumorigenic environment. Hence, germline genomic information could be used for developing non-invasive genomic tests for predicting patients’ outcomes (or drug response) in breast cancer, other cancer types and even other complex diseases.

## Introduction

Cancer is a process of asexual evolution driven by genomic alterations. A single normal cell randomly acquires a series of mutations that allows it to proliferate and to be transformed into a cancer cell (i.e., founding clone) that then **i**nitiates tumor progression and recurrence. In general, cancer recurrence and metastasis are the result of the interactions of multiple mutated genes. New somatic mutations arise and are selected if they confer a selective fitness advantage (e.g., proliferation, survival, etc.) to a founding clone in the context of a pre-existing genomic landscape (i.e., germline genetic variants). Hence, pre-existing germline genetic variants provide a profound constraint on the evolution of tumor founding clones and subclones, and therefore, have a contingent effect on the genetic makeup of tumor and presumably patient outcomes. Family history remains one of the major risk factors that contribute to cancer and recent studies have identified several genes whose germline mutations are associated with cancer. For example, patients suffering from Li-Fraumeni syndrome have an almost 100% chance of developing a wide range of malignancies before the age of 70. Most patients carry a missing or damaged *p53* gene, a tumor suppressor whose activity is impaired in almost 50% of all cancers. Other cancer-predisposition genes include *BRCA1* and *BRCA2*^1,2^, which are associated with breast and ovarian cancer, *PTEN*^*3*^, whose mutation results in Cowden syndrome, *APC*, which is linked to familial adenomatous polyposis^4^ and the Retinoblastoma gene *RB1*^*5*^. Two distinct types of multiple endocrine neoplasias are associated with the *RET* and *MEN1*^6^ genes while *VHL* alterations result in kidney and other types of cancer^7^. Finally, Lynch syndrome, a form of colorectal cancer, is linked to *MSH2, MLH1, MSH6, PMS2,* and *EPCAM*^8^. Genetic tests based on these highly-penetrant gene mutations have shown their usefulness, but they can explain only a small fraction (5-10%) of patients. Most neoplasms arise and are modulated by the interactions of multiple genes and there is a great diversity of genetic alterations even within tumors of the same subtypes.

Thus far, it is unclear to what extent germline genetic variants affect tumorigenesis, tumor evolution and even clinical outcome. We have previously shown that tumor founding clone mutations are able to predict tumor recurrence^9^. Here, we reasoned that the collective impact of germline genetic variants in cancer patients might largely determine tumorigenesis, evolution and even clinical outcomes. That is, germline genetic variants act in combination with newly acquired somatic mutations to modulate tumorigenesis and tumor recurrence. The combination of germline variants and somatic mutations of each patient predispose to the specific activation of biological/signaling pathways (even phenotypes) that directly impact clinical outcomes of cancer patients. Therefore, the germline genomic landscape of cancer patients might predict disease progression. Yet, thus far, clinical outcome predictions using cancer germline genomic information have been limited to only a few cancer types, or to a limited number of gene^1,2,3,4,5,6,7,8^. The increasing availability of genome sequencing data provide opportunities to develop predictive models that can translate these complex genomic alterations into clinical use.

In this study, we showed that the collective germline genetic variants of breast cancer patients predict tumor recurrence by applying a recently developed method, eTumorMetastasis^9^, to 755 breast cancer patients. Further statistical analyses showed that the leukocyte gene expression levels and tumor-infiltrating leukocytes (TIL) fractions within tumors between the two predicted groups were significantly different. Germline mutated variants associated with tumor recurrence likely impair the adaptive immune response functions of affected individuals, increasing the susceptibility to relapse. These results highlight the important role of germline genetic variants in tumor evolution and recurrence.

## Results

### Germline genetic variants predict breast cancer recurrence

To examine if germline genetic variants are able to predict tumor recurrence, we used whole-exome sequencing data (i.e., from the NCI Genomic Data Commons, GDC) of healthy tissues from 755 ER+ breast patients by applying our recently developed method, eTumorMetastasis^9^. ER+ subtype represents ~70% of breast cancer patients, thus, in this study, we used only patient data from this subtype. The demographic table of the breast cancer cohort is represented in Table 1.

**Table 1.**
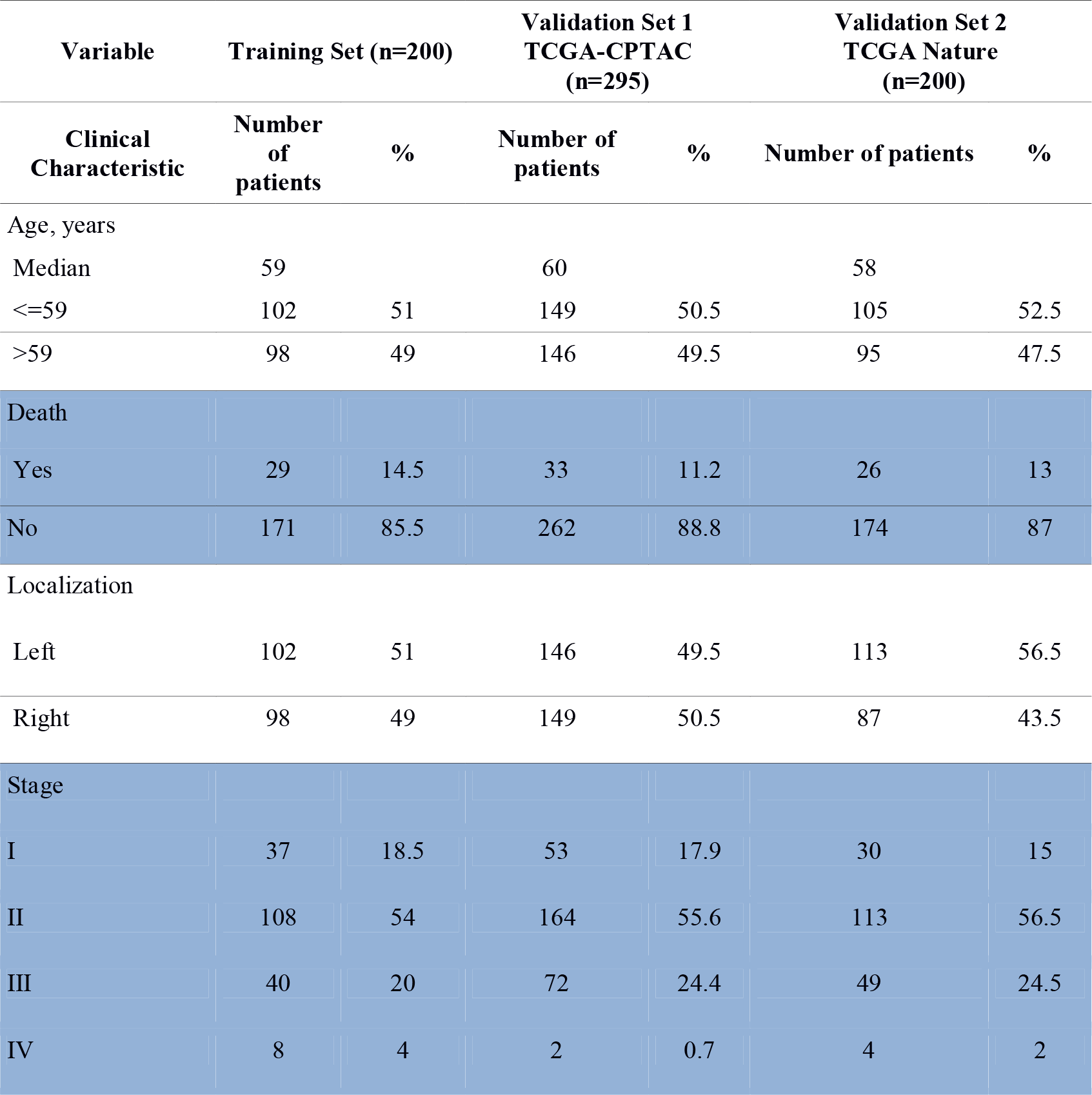
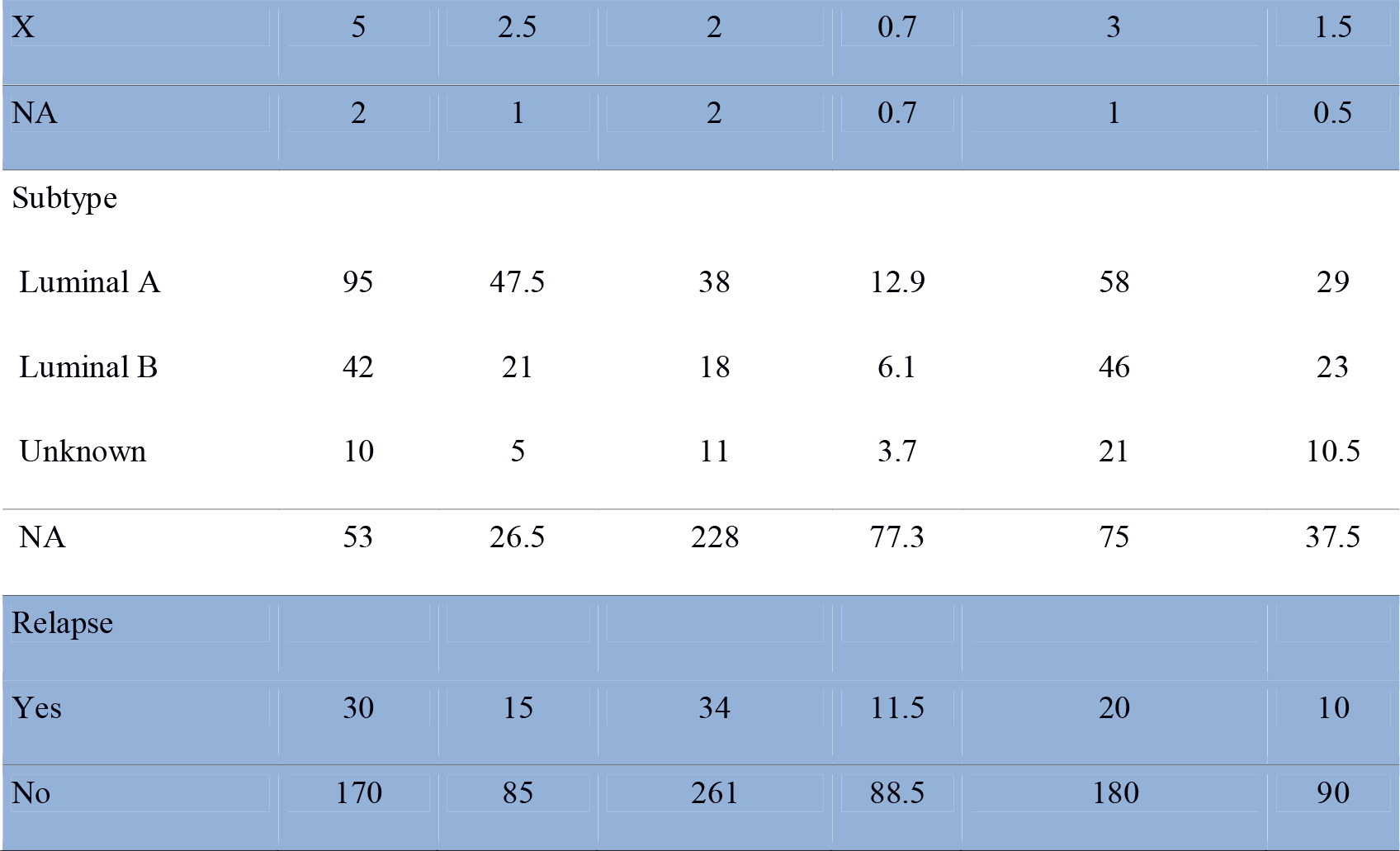
Demographic and clinical characteristics for ER+ breast cancer samples.

We hypothesized that somatic mutations are evolutionary selected to work with the pre-existing germline genetic variants to initiate tumorigenesis and recurrence. This is the underlying concept of eTumorMetastasis. In turn, the model infers that pre-existing germline genetic variants of cancer patients have predictive power for recurrence and clinical outcomes. eTumorMetastasis contains 3 main components: (1) a network-based approach^11,12^ to transform functionally genetic variants information on a cancer type specific signaling network; (2) identifying biomarkers via our previously developed method, MSS (Multiple Survival Screening)^13^ and (3) a better predictive power using our previously developed method by combining biomarkers^14^. The detailed procedure of eTumorMetastasis and network construction were described previously^9^. To apply eTumorMetastasis, briefly we first annotated the functional variants using the germline whole-exome sequencing data of each breast cancer patient (see Methods and Supplementary Methods), constructed an ER+ breast cancer-specific molecular network for recurrence (see Methods), and then mapped the functional germline-variant genes on the recurrence network. Finally, gene signatures (i.e., biomarkers) were obtained using MSS and eTumorMetastasis.

We used the germline genomic information of 200 ER+ breast cancer samples (i.e., training samples) to identify gene signatures (i.e., because eTumorMetastasis identifies network-based gene signatures, we called the gene signatures Network Operational Signatures or NOG signatures), which could distinguish recurred and non-recurred breast tumors. By applying eTumorMetastasis to the germline genomes of 200 patients, we identified 18 NOG signatures (Supplementary Tables 1 and 2) for ER+ breast cancer. Each NOG contains 30 genes and represents a cancer hallmark such as apoptosis, cell proliferation, metastasis, and so on. We have previously shown that multiple gene signatures representing distinct cancer hallmarks could be identified from one training cohort^13^. Furthermore, ensemble-based prediction using multiple gene signatures representing distinct cancer hallmarks significantly improved prediction performance^14^. Thus, we used the 18 NOG gene signatures to construct a NOG_CSS (i.e., NOG-based Combinatory Signature Set) using a testing set of 60 samples based on the method we developed previously^14^. We then used the NOG_CSS to predict the prognosis of ER+ breast cancer patients. As shown in Figure 1 and Table 2, we demonstrated that the germline-derived NOG_CSS significantly distinguished recurred and non-recurred breast tumors in two validations sets: 200 (ER+ Nature-Set, P=1.4×10^−2^), 295 (ER+ TCGA-CPTAC independent set, P=1.4×10^−3^). These results suggest that germline genetic variants are significantly correlated with tumor recurrence and support our hypothesis that the original germline genomic landscape of a cancer patient has a significant impact on clinical outcome.

**Figure 1.**
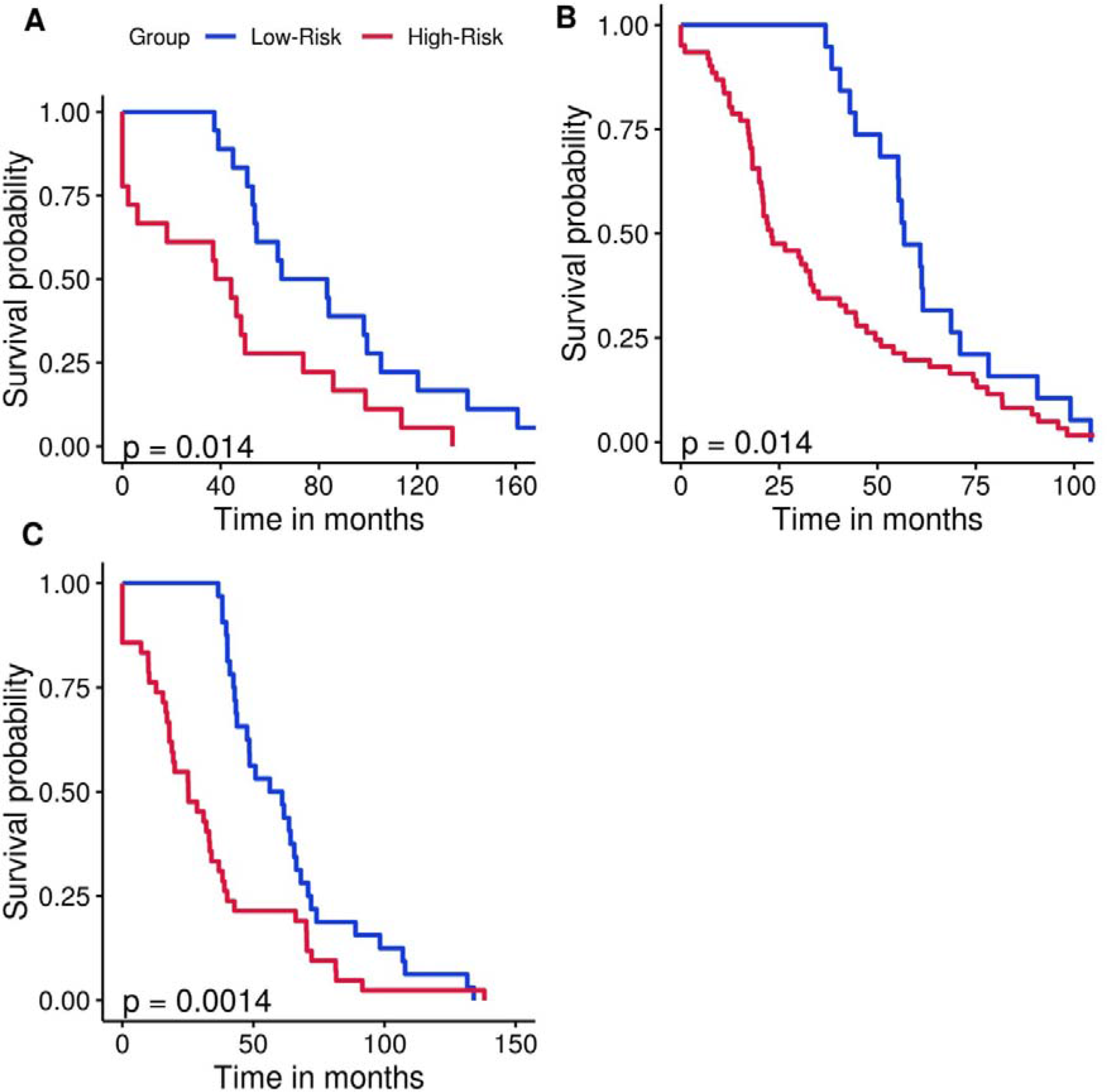
Kaplan–Meier curves of the risk groups for breast cancer patients with 10-year disease-free survival predicted by the NOG_CSS sets Samples without DFS time or who couldn’t be predicted were removed. NOG_CSS sets derived from germline mutations in (**A**) the training set, (**B**) the validation set, TCGA-Nature and (**C**) the validation set, TCGA-CPTAC. Blue and red curves represent low-and high-risk groups, respectively. P-values were obtained from the χ^2^-test.

**Table 2.** Prediction accuracy and recall rate for validation sets for breast cancer using the NOG_CSS sets derived from germline mutations.

| Dataset | Number of samples | Low-Risk |  | High-risk |  |
| --- | --- | --- | --- | --- | --- |
|  |  | Accuracy (%)<sup>*</sup> | Recall (%)<sup>†</sup> | Accuracy (%)<sup>**</sup> | Recall (%)<sup>††</sup> |
| Training Set | 200 | 93.8 | 26.5 | 27.5 | 36.7 |
| TCGA-Nature | 200 | 94.9 | 31.1 | 10.4 | 65.0 |
| TCGA-CPTAC | 295 | 93.5 | 38.7 | 21.0 | 50.0 |

Notes:
<sup>\*</sup>Percentage of non-recurred (i.e., non-metastatic) samples in the predicted low-risk group.
<sup>†</sup>Percentage of the predicted low-risk samples from the non-recurred group.
<sup>\*\*</sup>Percentage of recurred (i.e., metastatic) samples in the predicted high-risk group.
<sup>††</sup>Percentage of the predicted high-risk samples from the recurred group.

As a proof of concept and to further demonstrate the constraint given by germline variants onto the tumor development, we used the NOG_CSS and the gene expression of normal tissue of 72 breast cancer patients to predict patients relapse risk (See Methods for details). Samples were assigned in the training or validation set previously defined. The results of this prediction can be found in Table 3. Accuracy for low-risk samples was similar to germline variants predictions (88.9% compared to 94.9%) suggesting that the impact from germline variants is also reflected in gene expression and correlates with our hypothesis that gene expression and tumor development are affected directly from germline predispositions. Strikingly, the accuracy obtained for high-risk samples with gene expression data was much better than what we obtained using germline variants (66.7% compared to 21.0%), suggesting that gene expression is a better predictor of recurrence for high-risk patients or that high-risk patients might possess a more complex somatic landscape not captured solely by germline mutations.

**Table 3.** Prediction accuracy and recall rate for validation samples for breast cancer using the NOG_CSS sets derived from gene expression of normal tissue.

| Dataset | Number of samples | Low-Risk |  | High-risk |  |
| --- | --- | --- | --- | --- | --- |
|  |  | Accuracy (%)<sup>*</sup> | Recall (%)<sup>†</sup> | Accuracy (%)<sup>**</sup> | Recall (%)<sup>††</sup> |
| TCGA-Validation | 49 | 88.9 | 48.5 | 66.7 | 62.5 |

Notes:
<sup>\*</sup>Percentage of non-recurred (i.e., non-metastatic) samples in the predicted low-risk group.
<sup>†</sup>Percentage of the predicted low-risk samples from the non-recurred group.
<sup>\*\*</sup>Percentage of recurred (i.e., metastatic) samples in the predicted high-risk group.
<sup>††</sup>Percentage of the predicted high-risk samples from the recurred group.

To compare the prediction performance of the NOG_CSS with clinical factors, we conducted relapse-free survival analysis of clinical factors using Cox proportional hazards regression model. The best p-value (i.e., P=1.0×10^−2^., log-rank test) using covariate models (Supplementary Table 3) was not better than the one derived from the germline NOG_CSS (P=1.4×10^−3^). These results suggest that gene signatures derived from germline genomic information have a better predictive performance than clinical factors.

Finally, we also assessed the number of functional germline genetic variants in all genes or genes specifically expressed in leukocytes as well as the number of genes harboring germline genetic variants for both predicted risk group. Student T-tests revealed a significant difference for all the comparisons (1.29×10^−13^, 8.24×10^−16^ and 1.14×10^−5^, respectively) with functional germline genetic variants in leukocyte expressed genes being the most indicative distinction. All distributions are highlighted in Figure 2. A higher germline functional mutation count for high-risk group suggests once again that germline variants have a profound impact on tumor development and therefore, recurrence.

**Figure 2.**
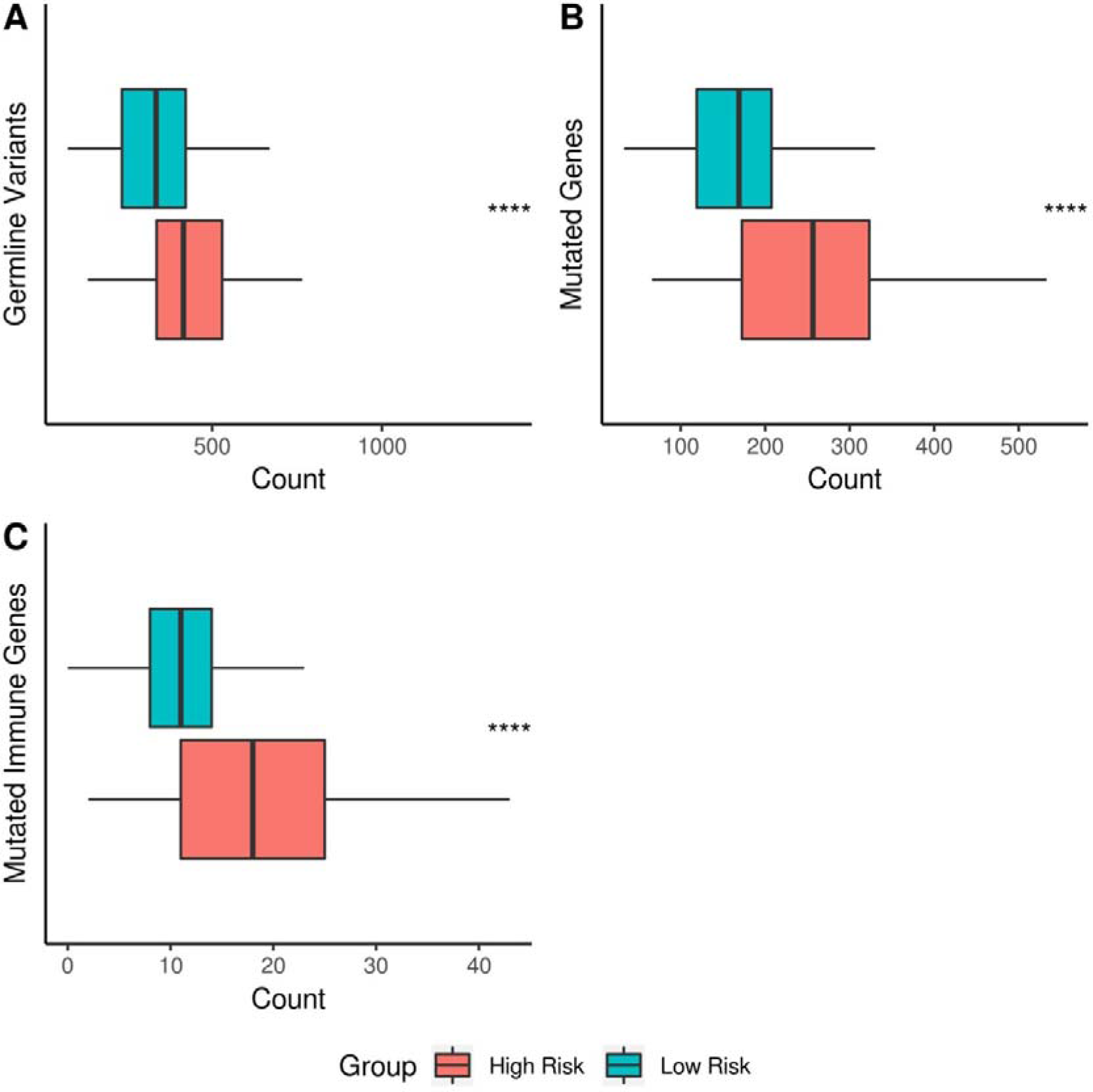
Boxplots comparison of functional germline mutated variants and genes for predicted risk groups. Samples who couldn’t be predicted were removed. **(A)** Functional germline variants **(B)** Functionally mutated genes **(C)** Germline mutated immune genes. P-Values were obtained from Student’s t-test. P-Value significance: **** < 0.0001.

### Predictive germline genetic variants potentially impair both the innate and adaptive immune system of breast cancer patients

To further understand why germline genomic landscapes of cancer patients are predictive for tumor recurrence, we ran enrichment analyses for genes present in the NOG signatures of breast cancers using DAVID^15^. Interestingly, most genes were enriched in immune-or cell proliferation-related biological pathways and Gene Ontology (GO) terms (Supplementary Tables 5).

Thus, we hypothesized that recurred patients have more functionally inherited variants in immune system related genes than non-recurred patients. To test this hypothesis, we compared gene expression for leukocytes metagenes between predicted recurred and non-recurred patients from tumor transcriptomes. The leukocytes metagene list was obtained from a recent study^16^. Student’s T-tests between both groups revealed a significant difference for myeloid-derived suppressor cells (MDSCs), effector memory CD8 T cells (E-Memory CD8+ T cells), activated dendritic cells (DCs+), activated CD8 T cells (CD8 T cells+), T follicular helper cells (Tfh), monocytes (Monos), memory B cells and activated B cells (B cells+)(P=1.99×10^−3^, P=4.03×10^−3^, P=6.67×10^−3^, P=2.10×10^−2^, P=2.30×10^−2^, P=3.78×10^−2^, P=4.37×10^−2^ and P=4.46×10^−2^, respectively). To a similar extent, we also analyzed TILs fractions to see if these were different between predicted groups (CIBERSORT LM22, see Methods)^16,17^. Student’s T-tests revealed a significant difference in TILs fractions for gamma delta T cells (γδ T cells), resting natural killer cells (NK cells-), resting mast cells (MCs-) and CD8+ T cells (P=3.14×10^−2^, P=4.29×10^−2^, P=4.97×10^−2^, P=8.21×10^−3^, respectively). A better representation of leukocytes gene expression and TILs fractions between the predicted groups are shown in Figure 3 and the complete abbreviation lists can be found in Supplementary Tables 6-7. Overall, these results suggest that germline genetic variants of cancer patients could directly influence gene expression and alter immune system functions, cell division and the immune tumor microenvironment (TME). Modulation of these pathways would then affect recurrence and patient outcome.

**Figure 3.**
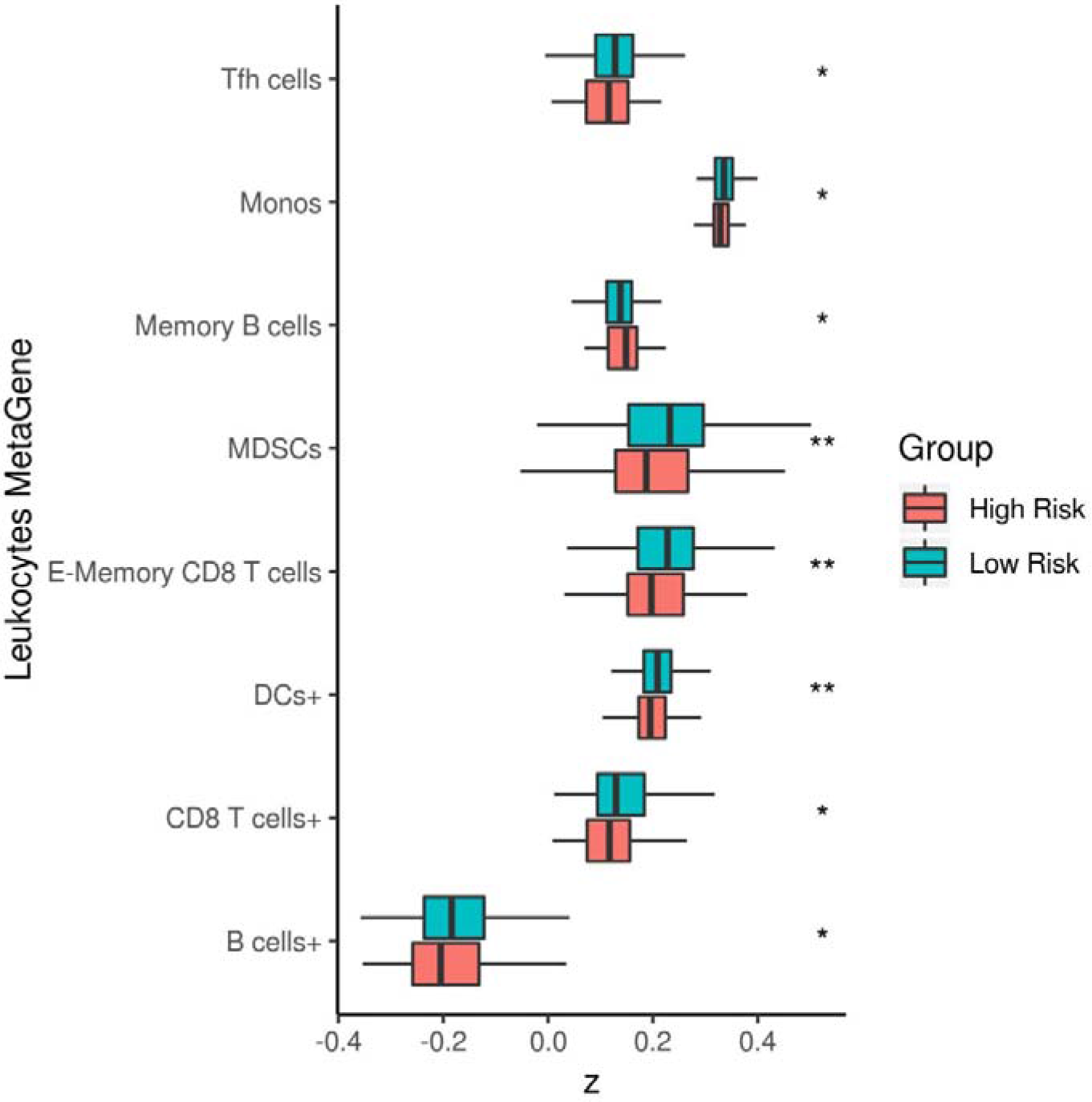
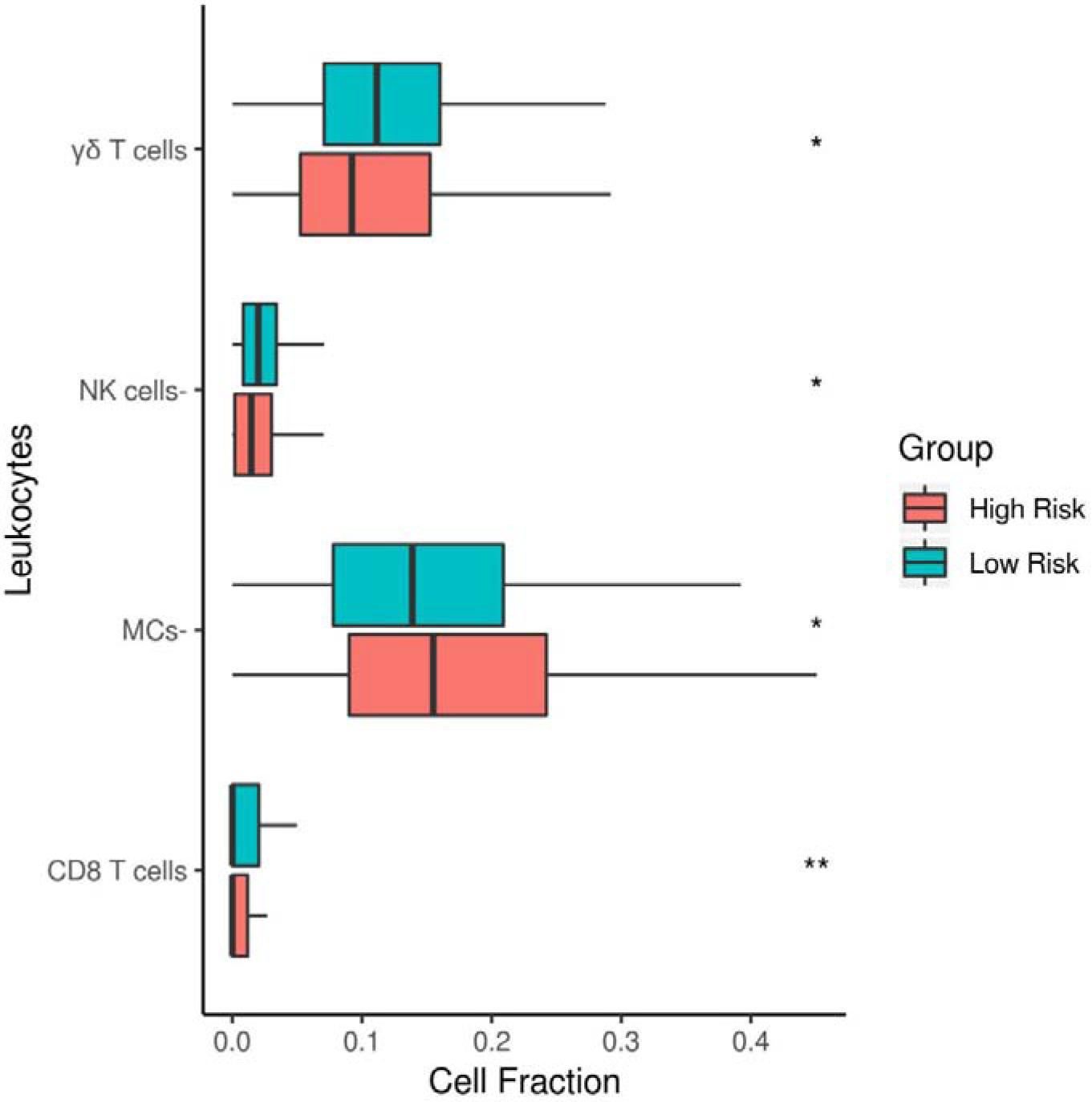
Boxplots comparison of leukocytes metagenes and cell fractions for predicted risk groups Samples who couldn’t be predicted were removed. For a complete analysis, see Supplementary Figure 1. (**A**) Leukocytes gene expression (**B**) Leukocytes cell fraction. P-Values were obtained from Student’s t-test. P-Value significance: * < 0.05, ** < 0.01.

To further investigate the predictive power of variants in leukocytes-expressed genes, we re-ran eTumorMetastasis^9^ pipeline using only functional germline variants in leukocytes-expressed genes (Supplementary Methods). Interestingly, we weren’t able to obtain enough significant gene sets to extract a gene signature proposing that leukocytes variants only provides partial information and that the complete germline mutational landscape is more representative.

## Discussion

With the recent advances in new generation sequencing, genetic testing and patients monitoring has increased drastically. Yet, it remains a challenge to stratify patients in order to provide an optimal treatment route. Here, we developed a risk classification method using germline genomic variants to predict clinical outcomes and demonstrate that these germline variants shape tumor evolution and recurrence. The enrichment analysis of the NOG signatures derived from germline genetic variants suggest that recurred patients differently regulate signaling pathways associated with immune responses (such as inflammation and cell adhesion).

Breast cancer patients with no lymph node involvement often undergo unnecessary adjuvant chemotherapy treatment (70-80% of patients). In fact, toxic therapies are given to most women with early-stage breast cancer from which 60–75% will not receive any benefit, but instead will experience only side effects^19^. Therefore, biomarkers identification to accurately stratify low-risk breast cancer patients who will not benefit from adjuvant chemotherapy is essential. We showed that germline variants enabled to identify low-risk patients with an accuracy as high as 94.9%, suggesting that inherited variants of breast cancer patients are useful in clinical applications.

Comparison of germline variants and affected genes between the two predicted groups indicates that these variants are predisposing to cancer. Potentially, a significantly higher number of functional variants could lead to a greater number of impaired proteins which would create an imbalance in signaling pathways, favoring tumor development and recurrence. As mentioned above, a functionally mutated gene does not always translate into an expressed loss-of-function protein but the difference between both groups suggest that nonetheless, the greater number of germline variants increases patient susceptibility to relapse. Moreover, we found that leukocyte genes harbored a greater number of germline genetic variants in the predicted high-risk group. These germline genetic variants likely impede the immune system, leading to a more favorable environment for tumor developments.

Our study suggests that tumor recurrence is predicted by the germline genomic landscapes of cancer patients. We found that germline variants in genes regulating cell division, immune cell infiltration and T cell activities are predominately predictive for tumor recurrence. More specifically, mutations in the antigen processing and presentation pathway could impair neoantigens presentation at the surface of cancer cells so that T cells are no longer able to recognize tumor cells, allowing them to evade immune detection. Furthermore, mutations in cell division process could introduce a higher number of somatic mutations during cell division directly promoting tumor development. Activation of Wnt pathway can also block the infiltration of immune cells within tumors^20^. TILs fractions analyses also reveal strong correlation with germline prediction and differential expression in MDSCs, CD8+ T cells, DCs, Tfh cells, monocytes and B cells (Fig. 3A). Aside from memory B cells, all other TILs were enriched in the predicted low-risk group. B cells have been shown to secrete pro-tumorigenic factors (e.g., angiogenesis, tumor growth) and also to inhibit the anti-tumor immune response via cytokines^21–23^. DCs are well known for their role in antigen presentations and in initiating an adaptive immune response^24^. Tfh cells have been shown to favor an adaptive immune response via the B cell chemoattractant CXLC13 in breast cancer^25^. Along with E-memory CD4 T cells, E-memory CD8 T cells possess a key role in the immune response and tumor infiltration. Patient survival has been directly correlated with CD8 T cells infiltration. Multiple mechanisms are used by cancer cells to escape immune responses such as altering cytokines and chemokines attraction to create a non-inflammatory environment which, in turn, inhibits T cells infiltration^26,27^. Monocytes and MDSCs have largely been associated with tumor recurrence in the literature. Monocytes differentiation into tumor-associated macrophages (TAMs) promotes anti-immunity signals such as angiogenesis and growth factors resulting in a TME favoring cancer cell proliferation. However, there have been some reports indicating that a nonclassical monocyte subtype, patrolling monocytes, reduces tumor recurrence by recruiting NK cells^28,29^. Monocytes can also differentiate into pro-inflammatory M1 macrophages aiding the adaptive immune response. A recent study has also shown that TNFα secreted by T cells induces emergency myelopoiesis resulting in an increase in MDSCs in mice^30^. TNFα secretion by T cells could be a regulation mechanism induced by the adaptive immune response once a certain concentration of T cells has infiltrated the tumor. This point could explain the higher expression numbers for MDSCs in predicted low-risk samples.

A significant difference was also seen in TILs cell fractions of γδ T cells, CD8 T cells, NK cells-and MCs (Fig. 3B) between both predicted groups. As mentioned above, CD8 T cell tumor infiltration is crucial for an optimal immune response; these cells were present in greater numbers in the predicted low-risk group. Gamma delta T cells (γδ T cells) are known to have dual effects, capable of exerting both pro- or anti-tumor response depending on their subtype^31^. γδT1, γδT-APC and γδTfh subtypes all possess antitumor activity such as secreting chemoattracting chemokines (i.e. CXLC13), antigen presentation and antibody-dependent cell-mediated cytotoxicity towards cancer cells^32^. Tumor-infiltrating mast cells (MCs) can either emit pro-or anti-tumor signals. Specifically, in breast cancer, MCs are linked with pro-angiogenic factors such as inflammation^33,34^ reflecting a higher MCs count in the predicted high-risk group. Finally, NK cells have cytotoxic abilities and a greater number in tumors is indicative of a good prognosis^35,36^. Overall, germline genetic variants identified in the high-risk group could directly modulate gene expression in immune cells, resulting in a weakened adaptive immune response.

Our understanding of the biology mediating recurrence is limited. Germline variants of cancer patients could affect the activity of the immune system in TMEs. For example, germline-encoded receptor variants were shown to trigger innate immune response in cancer patients^37^. In addition, lung cancer patients with a germline mutation in Nrf2 have a good prognosis because these variants regulate the inflammatory status and redox balance of the hematopoietic and immune systems of cancer patients^38^. In prostate cancer, patients with a germline variant of the ASPN D locus are associated with poorer outcomes^39^. These studies, including our own, highlight the impacts of germline variants on tumor recurrence and provide a rationale to further study the effect of germline genomic landscapes on clinical outcomes of carcinogenesis.

Good accuracy obtained using normal tissue RNA prediction shows that germline variants directly influence gene expression and consequently, tumor development. A higher accuracy for the high-risk group also highlights that gene expression holds a better predictive power than genome sequencing. These results are not surprising considering that gene expression integrates more information than gene-coding mutations alone (e.g, gene regulation). Even the most damaging functional mutation in a gene not expressed would have no impact on the phenotype. However, gene expression usually requires a biopsy. Therefore, exome-sequencing of normal tissue provides a much more convenient and less invasive method for clinical purposes. We also note that this analysis suffers from a small sample size and should be further explored in the future.

In all, these results suggest that germline variants potentially alter the immune system and the immune TME which in turn stimulate tumor recurrence and ultimately, affect patient outcome. Traditionally, germline variants have been largely ignored in the cancer genomic community; for example, most of the cancer genomic studies including the GDC and The Cancer Genome Atlas (TCGA) have often focused only on somatic mutations while germline mutations were filtered out before formal analysis of tumor genome sequencing data. The demonstration that germline exome sequencing data can predict cancer patients’ outcomes suggests that non-invasive genomic tests of cancer patients could be devised to determine cancer prognosis and inform treatment decisions. Genome-wide germline genetic variants can be easily identified by genome/whole-exome sequencing of liquid biopsies such as blood or saliva samples. Prognostic prediction using a patient’s germline genomic landscape opens up the possibility of assessing cancer patients’ risk of recurrence in a non-invasive manner, which allows for a better forecasting of cancer recurrence in a quick, convenient and minimally invasive manner.

## Methods

### Exome data processing

We obtained whole-exome sequencing data of breast cancers from the GDC: ER+ breast cancer, a training set of 200 samples, a testing set of 60 samples and two independent validation sets of 200 and 295 samples (TCGA-Nature and TCGA-CPTAC, respectively, Supplementary Table 4). Raw sequence reads from healthy samples of cancer patients were processed using GATK^40^ pipeline and the method described previously^9^. Variant calling was then performed using Varscan2^41^.

### Transcriptome data processing

Normal tissue RNA-seq is less accessible on the GDC than tumor RNA-seq data. Out of 755 samples in our dataset, we were only able to find 72 samples from which normal tissue RNA-seq was available. FPKM values for each sample were downloaded and then normalized using z-score normalization. Each sample was then assigned to our previously defined training and validation set (23 and 49, respectively).

### Germline variant identification

To determine germline variants, we used variant allele frequencies (VAFs) between the tumor and healthy samples. We defined homozygous germline variants if the VAF in the healthy samples was >=90. For heterozygous germline variants, we used the VAF cutoffs between 45 and 65% in healthy samples. Only germline functional variants were retained for downstream analysis.

### Germline NOG signature identification

To identify NOG signatures using the functional mutated genes of breast cancer patients’ germline genomes, we followed the eTumorMetastasis^9^ method. Briefly, we constructed a breast-specific recurrence network based on the methods described previously^9^. For each patient, we used its germline functionally mutated genes as seeds on the breast cancer-specific recurrence network to perform network propagation and then identify NOG signatures.

### Transcriptomic normal tissue prediction

Like mentioned above, each sample was assigned to our previously defined training and validation set (23 and 49, respectively). Accuracy and recall rate were obtained using a similar approach than with the eTumorMetastasis^9^ method. For all 18 NOG signatures previously identified with genome sequencing, we calculated centroids values for each gene between both groups (high-and low-risk) in the training set. In this case, centroids values were obtained from gene expression values instead of network propagation scores. For each sample in our validation set, each sample was classified to its specific group based on Pearson correlation with centroids from both groups. We built a NOG_CSS using the same cutoffs obtained from genome sequencing. Prediction accuracy and recall rate for validation samples can be found in Table 3.

### Leukocytes metagene expression and cell fractions

Leukocytes metagene expression derived from tumor RNA-seq data were obtained from The Cancer Immunome Atlas (TCIA)^16^ and were applied z-score normalization. In total, scores for 29 leukocytes metagene were downloaded. Leukocytes cell fractions were also downloaded from TCIA for all 755 breast cancer samples. CIBERSORT^17^ signature of 22 leukocytes cells was used (LM22).

## Author contributions

Manuscript draft: JSM, AN, RM, SD, EW.

Conceptualization and implementation: JSM, CT, EW.

Data acquisition: JSM, JZ, PH, ZM.

Data analysis: JSM, CT.

Data interpretation: JSM, CT, RM, EW.

