## Supplementary Files for "Germline genetic variants associated with leukocyte-genes predict tumor recurrence in breast cancer patients"

**Supplementary Table 1 List of genes of network operational signatures derived from breast cancer germline mutations**

| **Apop1** | **Apop2** | **Apop3** | **CCycle1** | **CCycle2** | **Ccycle3** | **Cell Adh1** | **Cell Adh2** | **CellAdh3** | **Cytosk1** | **Cytosk2** | **Cytosk3** | **Imm Res1** | **Imm Res2** | **ImmRes3** | **Prolif1** | **Prolif2** | **Prolif3** |
| --- | --- | --- | --- | --- | --- | --- | --- | --- | --- | --- | --- | --- | --- | --- | --- | --- | --- |
| BRCA1 | APOE | ANXA1 | ACVR1 | BRSK1 | BRCA1 | ADAM17 | ADAM17 | ADAM17 | ACTA2 | ACTRT1 | ACTRT1 | CD19 | BCAP31 | B2M | AGT | ADRA1B | ANXA1 |
| CASP8 | AVEN | BCAP31 | ANXA1 | CCNB1 | C13orf34 | AMBN | BRCA1 | CAV1 | ACTR2 | APOE | AKAP9 | CD4 | CCL4 | C3 | AMBN | BIRC2 | BOK |
| CDK11A | BIRC2 | BFAR | ATR | CCNB3 | CCNB1 | BAI1 | CCL4 | CD209 | ACTRT1 | BBS4 | ALMS1 | CD81 | CCR5 | CCL23 | ASCL1 | BMPR2 | CD81 |
| DOCK1 | BRCA1 | BRCA1 | BRD7 | CDK11A | CCNB3 | CD209 | CD209 | COL11A2 | ASAP1 | CASP8 | CASP8 | CHIA | CD160 | CCL3 | CD81 | BRCA1 | CITED1 |
| EP300 | CDK11A | CDK11A | BRSK1 | CHFR | CDC25B | CD4 | COL6A3 | COL6A3 | BRSK1 | CDK5RAP2 | CYTH2 | CIITA | CD19 | CD19 | CITED1 | CD160 | CITED2 |
| ESPL1 | DAPK1 | CITED2 | CASP8AP2 | CLASP1 | CDCA5 | CD58 | CTGF | CTNND1 | CASP8 | CENPJ | DAPK3 | HCST | CD81 | CD81 | CSF1R | CD81 | CKS1B |
| FGF2 | ESPL1 | ESPL1 | CCNB1 | DCTN2 | CDK11A | CDH17 | DDR1 | EPHA3 | CCNB1 | CEP72 | DIAPH2 | HLA-DMB | CIITA | CIITA | DBH | CDK9 | CTF1 |
| GPX1 | GSN | FAIM3 | CCNB3 | DDIT3 | CHFR | CDH4 | EZR | FAT1 | CDC25B | CEP78 | GSG2 | HLA-DOA | FASLG | ENPP2 | GHRL | CITED1 | DAB2 |
| GSN | GULP1 | FBXO7 | CDK11A | E2F6 | FBXO31 | COL5A1 | F11R | FLOT2 | DAPK3 | CKAP5 | GSN | HLA-DOB | HCST | HCST | GNB1 | DCTN2 | DBH |
| HIPK1 | GZMB | GSN | CENPJ | GSG2 | KIF11 | COL6A3 | FAT1 | FN1 | GSN | CLASP1 | LSP1 | HLA-DPA1 | ICAM1 | ICAM2 | IGSF8 | EDNRA | FAS |
| HYAL1 | GZMH | HIPK1 | CHAF1B | KIF11 | NASP | DPP4 | FLOT2 | IBSP | KIF11 | DAPK1 | MAP2K1 | HLA-DQA2 | ICAM2 | ICAM3 | LTK | ENG | GHRL |
| IKBKB | HIPK1 | IER3 | CHFR | MAPK7 | NCAPD2 | ENG | FN1 | ITGAM | LSP1 | DAPK3 | NCAPG | HLA-DRB3 | ICAM3 | IFITM1 | MDM4 | GHRL | GNB1 |
| IL17A | IKBKB | IKBKB | EP300 | MRE11A | NCAPG | FAT1 | ICAM1 | LAYN | NCAPG | DCTN2 | NF1 | HLA-DRB4 | IFITM1 | IL17A | MMP7 | GNB1 | HOXB4 |
| IL7 | IL17A | IL17A | FBXO31 | NASP | NOTCH1 | FLOT2 | IGFALS | LEF1 | NCAPH | DCTN3 | NPM1 | HLA-DRB5 | IL15 | IL21 | NASP | IL2RA | LTK |
| MAP2K7 | IL7 | IL7 | FOXN2 | NCAPD3 | OFD1 | FN1 | ITGA4 | LIMS1 | NF1 | DIAPH1 | PCM1 | ICAM2 | IL17A | IL7 | NCF1 | INHA | MPL |
| NEFH | LTK | LTBR | KIF11 | NCAPG | PIN1 | ITGAL | ITGAV | MAP2K1 | NPM1 | EZR | PCNT | ICAM3 | IL7 | ITGA4 | PIM2 | MRE11A | PAK7 |
| PIM2 | PLEC | MDM4 | LZTS1 | NOTCH1 | PINX1 | LEF1 | JAM3 | MAP2K2 | NUCB1 | GSN | PDPK1 | ICAM4 | IL9 | ITGAL | PLAU | PLAU | PIN1 |
| PSMD5 | PSME1 | MUC1 | MAP2K1 | OFD1 | PLK1 | LY9 | LEF1 | MLLT4 | OFD1 | GTF2F2 | PFN1 | IFITM1 | ITGAL | KLRC1 | PLK1 | PLK1 | PLAU |
| PSME1 | PSME4 | PCNT | MDC1 | P2RY2 | RAD21 | MAP2K2 | MAP2K2 | MOG | PLK1 | LSP1 | PLEKHG6 | IL17A | KLRD1 | NOTCH1 | PPP1R9B | PPP1R9B | PPP1R9B |
| PSME4 | RASSF5 | PIM2 | MTBP | PINX1 | RBBP8 | MOG | MDC1 | MUC4 | PTPN13 | NCAPG | PLK1 | IL18 | MAP3K14 | PDCD1LG2 | RAP1B | PTEN | SAT1 |
| PTK2B | SEMA3A | PSME1 | NASP | PPP1CA | REC8 | MUC4 | MOG | NFASC | RASSF5 | NF1 | PTK2B | IL2RG | MICB | RAET1G | SAT1 | SAT1 | SGK1 |
| RAD21 | SERPINB9 | PSME4 | NCAPG | RAD50 | RNF8 | MYH6 | MUC4 | NLGN4X | RPA1 | NPM1 | RASSF1 | IL7 | NCF4 | RAET1L | SCRIB | SRA1 | SOX9 |
| SEMA3A | SRA1 | SLK | NOTCH1 | RNF8 | RPA1 | NLGN4X | NLGN4X | NRP1 | S100A9 | PLK1 | RASSF5 | ITGAL | NOTCH1 | SPN | SPHK2 | TACC3 | TCF7L2 |
| SMNDC1 | TGFBR2 | TGFB2 | OFD1 | RPA1 | RPS6KA1 | NPNT | NRP1 | OMD | SCNN1A | PSMD10 | RPA1 | NOTCH1 | RAET1E | TLR10 | TCF7L2 | TCF7L2 | TGFB2 |
| TGFBR2 | TNFRSF10B | TGFBR2 | PIM2 | RPS6KA1 | SMC1A | NRP1 | OMD | PAK1 | SORBS1 | RASSF5 | S100A9 | RAET1G | RAET1G | TNFRSF4 | TGFB2 | TGFB2 | TGFBR2 |
| TMEM173 | TNFSF10 | TNFRSF10B | PINX1 | SMC1B | SMC2 | OMD | PDPK1 | PKN2 | TESK2 | RPA1 | SORBS1 | RGS1 | SERPINB9 | TNFSF10 | TGFBR2 | TGFBR2 | THPO |
| TNFRSF10B | TNFSF9 | TNFRSF25 | RNF8 | SMC3 | SMC3 | PTK2B | SORBS1 | PLEC | TPM1 | S100A9 | TPM1 | TLR10 | TLR10 | TNFSF15 | TNFSF8 | TNFSF9 | TNFSF9 |
| TNFSF10 | TP53BP2 | TNFSF10 | RPA1 | TGFB2 | SMC4 | S1PR1 | TENC1 | SORBS1 | TSSK2 | SORBS1 | TUBA4B | TNFSF10 | TNFSF10 | ULBP1 | TNFSF9 | UBE2L3 | UBE2L3 |
| TREX1 | TP63 | TREX1 | RPS6KA1 | TGFB3 | STAG1 | SORBS1 | TPPP | TNS1 | UBXN6 | TPM1 | UNC119 | ULBP2 | TNFSF9 | ULBP3 | UBE2L3 | UCHL1 | UCHL1 |
| YARS | TREX1 | VHL | SMC3 | USP2 | STAG2 | TPPP | VCAM1 | TPPP | UNC119 | UNC119 | WIPF1 | ULBP3 | ULBP3 | VCAM1 | UCHL1 | DBH | ZEB1 |

Abbreviations: Apop, Apoptosis; CCycle, Cell Cycle; CellAdh, Cell Adhesion; Cytosk, Cytoskeleton, ImmRes, Immune Response; Prolif, Cell Proliferation.

**Supplementary Table 2 Prediction accuracy and recall rate for the gene signatures derived from germline mutations of breast cancer tumors**

| **Dataset** | **Number of samples** | **Cancer Hallmark** | **Low-risk** | | **High-risk** | |
| --- | --- | --- | --- | --- | --- | --- |
|  |  |  | **Accuracy (%)*** | **Recall (%)^†^** | **Accuracy (%)**** | **Recall (%)^††^** |
| Training | 200 | Apoptosis 1 | 94.8 | 83.3 | 24.0 | 53.5 |
|  |  | Apoptosis 2 | 94.9 | 83.3 | 24.5 | 54.7 |
|  |  | Apoptosis 3 | 91.7 | 73.3 | 21.2 | 51.8 |
|  |  | Cell Cycle 1 | 92.3 | 76.7 | 21.1 | 49.4 |
|  |  | Cell Cycle 2 | 91.4 | 70.0 | 21.9 | 55.9 |
|  |  | Cell Cycle 3 | 91.1 | 70.0 | 21.2 | 54.1 |
|  |  | Cell Adhesion 1 | 90.1 | 66.7 | 20.2 | 53.5 |
|  |  | Cell Adhesion 2 | 84.4 | 53.3 | 14.6 | 44.7 |
|  |  | Cell Adhesion 3 | 93.7 | 80.0 | 22.9 | 52.4 |
|  |  | Cytoskeleton 1 | 86.7 | 56.7 | 16.7 | 50.0 |
|  |  | Cytoskeleton 2 | 77.0 | 23.3 | 7.0 | 45.3 |
|  |  | Cytoskeleton 3 | 78.7 | 36.7 | 9.9 | 41.2 |
|  |  | Immune Response 1 | 90.5 | 70.0 | 20.0 | 50.6 |
|  |  | Immune Response 2 | 87.9 | 60.0 | 17.8 | 51.2 |
|  |  | Immune Response 3 | 86.9 | 56.7 | 16.8 | 50.6 |
|  |  | Cell Proliferation 1 | 85.6 | 50.0 | 15.6 | 52.4 |
|  |  | Cell Proliferation 2 | 86.4 | 53.3 | 16.5 | 52.4 |
|  |  | Cell Proliferation 3 | 93.6 | 80.0 | 22.6 | 51.8 |
| TCGA-Nature | 200 | Apoptosis 1 | 89.4 | 42.2 | 9.6 | 42.2 |
|  |  | Apoptosis 2 | 93.8 | 50.6 | 13.6 | 70.0 |
|  |  | Apoptosis 3 | 88.8 | 48.3 | 8.8 | 45.0 |
|  |  | Cell Cycle 1 | 89.0 | 49.4 | 9.0 | 45.0 |
|  |  | Cell Cycle 2 | 90.6 | 48.3 | 10.6 | 55.0 |
|  |  | Cell Cycle 3 | 91.8 | 50.0 | 11.8 | 60.0 |
|  |  | Cell Adhesion 1 | 88.0 | 48.9 | 8.0 | 40.0 |
|  |  | Cell Adhesion 2 | 91.1 | 56.7 | 11.4 | 50.0 |
|  |  | Cell Adhesion 3 | 83.2 | 43.9 | 3.8 | 20.0 |
|  |  | Cytoskeleton 1 | 87.1 | 41.1 | 7.8 | 45.0 |
|  |  | Cytoskeleton 2 | 88.3 | 46.1 | 8.5 | 45.0 |
|  |  | Cytoskeleton 3 | 85.9 | 43.9 | 6.5 | 35.0 |
|  |  | Immune Response 1 | 90.4 | 41.7 | 10.3 | 60.0 |
|  |  | Immune Response 2 | 88.2 | 37.2 | 8.9 | 55.0 |
|  |  | Immune Response 3 | 88.3 | 37.8 | 8.9 | 55.0 |
|  |  | Cell Proliferation 1 | 87.0 | 44.4 | 7.4 | 40.0 |
|  |  | Cell Proliferation 2 | 93.1 | 52.2 | 13.1 | 65.0 |
|  |  | Cell Proliferation 3 | 92.2 | 52.8 | 12.4 | 60.0 |
| TCGA-CPTAC | 295 | Apoptosis 1 | 91.1 | 58.6 | 15.0 | 55.9 |
|  |  | Apoptosis 2 | 92.7 | 58.6 | 16.9 | 64.7 |
|  |  | Apoptosis 3 | 92.0 | 57.1 | 15.8 | 61.8 |
|  |  | Cell Cycle 1 | 90.1 | 52.5 | 13.3 | 55.9 |
|  |  | Cell Cycle 2 | 90.3 | 49.8 | 13.3 | 58.8 |
|  |  | Cell Cycle 3 | 89.2 | 57.1 | 12.5 | 47.1 |
|  |  | Cell Adhesion 1 | 87.2 | 49.4 | 10.2 | 44.1 |
|  |  | Cell Adhesion 2 | 86.5 | 51.3 | 9.3 | 38.2 |
|  |  | Cell Adhesion 3 | 85.9 | 53.6 | 8.3 | 32.4 |
|  |  | Cytoskeleton 1 | 87.9 | 58.6 | 10.7 | 38.2 |
|  |  | Cytoskeleton 2 | 91.1 | 54.8 | 14.5 | 58.8 |
|  |  | Cytoskeleton 3 | 91.4 | 57.1 | 15.2 | 58.8 |
|  |  | Immune Response 1 | 90.8 | 60.5 | 14.9 | 52.9 |
|  |  | Immune Response 2 | 91.4 | 60.9 | 15.7 | 55.9 |
|  |  | Immune Response 3 | 92.4 | 60.5 | 16.9 | 61.8 |
|  |  | Cell Proliferation 1 | 88.7 | 54.0 | 11.8 | 47.1 |
|  |  | Cell Proliferation 2 | 88.8 | 48.7 | 11.8 | 52.9 |
|  |  | Cell Proliferation 3 | 89.5 | 55.7 | 12.8 | 50.0 |

Notes:

*Percentage of non-recurred (i.e., non-metastatic) samples in the predicted low-risk group.

†Percentage of the predicted low-risk samples from the non-recurred group.

**Percentage of recurred (i.e., metastatic) samples in the predicted high-risk group.

††Percentage of the predicted high-risk samples from the recurred group.

**Supplementary Table 3 Cox proportional hazards regression model of uni- and multiple-factor for breast cancer**

| **Variable** | **P-Value** | **HR** | **95% CI** |
| --- | --- | --- | --- |
| Age | 0.13 | 1.006762 | 0.998-1.016 |
| Subtype, Luminal A v Luminal B | 0.04 | 1.32708 | 1.0194-1.728 |
| Subtype, Luminal A v Unknown | 0.82 | 0.95363 | 0.6335-1.436 |
| Subtype, Luminal B v Unknown | 0.15 | 1.3874 | 0.8933-2.155 |
| Localization, Left v Right | 0.03 | 0.7659 | 0.6014-0.9755 |
| Stage, I v II | 0.31 | 1.1703 | 0.8612-1.590 |
| Stage, I v III | 0.28 | 1.2240 | 0.8475-1.768 |
| Stage, I v IV | 0.06 | 0.2250 | 0.0622-1.045 |
| Stage, I v X | 0.31 | 0.5482 | 0.1715-1.752 |
| Subtype + Localization + Stage, Luminal A v Luminal B | 0.02 | 1.38377 | 1.05771-1.810 |
| Subtype + Localization + Stage, I v IV | 0.04 | 0.23098 | 0.05586-0.9551 |

Abbreviations: HR, hazard ratio; CI, Confidence Interval

**Supplementary Table 4 Sample filtering steps in breast cancer dataset**

| **Dataset** | **Clinical Information** | **Sequencing** | **Training Set** | **Testing Set** | **Validation Set 1 (TCGA-CPTAC)** | **Validation Set 2 (TCGA-Nature)** |
| --- | --- | --- | --- | --- | --- | --- |
| Breast | 1067 | 755 | 200 | 60 | 295 | 200 |

**Supplementary Table 5 Pathway enrichment analysis of network operational signature genes from germline mutations of breast cancer patients**

| **Category** | **Term** | **FDR PValue** |
| --- | --- | --- |
| GOTERM | Antigen processing and presentation | 8.38E-09 |
| KEYWORDS | Mitosis | 2.12E-08 |
| PATHWAY | Cytokine-cytokine receptor interaction | 3.01E-08 |
| GOTERM | Cell division | 1.99E-06 |
| GOTERM | Natural killer cell lectin-like receptor binding | 2.14E-06 |
| PATHWAY | Leishmaniasis | 7.55E-06 |
| GOTERM | T cell costimulation | 1.10E-05 |
| PATHWAY | Graft-versus-host disease | 1.43E-05 |
| GOTERM | Natural killer cell mediated cytotoxicity | 1.64E-05 |
| PATHWAY | Rheumatoid arthritis | 1.74E-05 |
| PATHWAY | Inflammatory bowel disease (IBD) | 1.84E-05 |
| GOTERM | MHC class II protein complex | 4.02E-05 |
| PATHWAY | Viral myocarditis | 4.40E-05 |
| PATHWAY | Allograft rejection | 4.87E-05 |
| PATHWAY | Intestinal immune network for IgA production | 5.15E-05 |
| PATHWAY | Negative regulation of extrinsic apoptotic signaling pathway via death domain receptors | 9.50E-05 |
| PATHWAY | Antigen processing and presentation | 1.60E-04 |
| GOTERM | Mitotic chromosome condensation | 1.62E-04 |
| PATHWAY | Type I diabetes mellitus | 1.82E-04 |
| PATHWAY | Staphylococcus aureus infection | 2.39E-04 |
| PATHWAY | Influenza A | 3.50E-04 |
| PATHWAY | Toxoplasmosis | 9.54E-04 |
| PATHWAY | Asthma | 0.00131511 |
| PATHWAY | Autoimmune thyroid disease | 0.00153788 |
| PATHWAY | Tuberculosis | 0.00210254 |
| KEYWORDS | Host-virus interaction | 0.0025258 |
| PATHWAY | HTLV-I infection | 0.00252675 |
| PATHWAY | FoxO signaling pathway | 0.00487651 |
| PATHWAY | Cell cycle | 0.00936092 |
| PATHWAY | Stimulatory C-type lectin receptor signaling pathway | 0.01846333 |
| PATHWAY | Negative regulation of canonical Wnt signaling pathway | 0.03468057 |
| KEYWORDS | Host cell receptor for virus entry | 0.03776077 |
| PATHWAY | Interferon-gamma-mediated signaling pathway | 0.04199796 |
| GOTERM | Positive regulation of inflammatory response | 0.05155117 |

**Supplementary Table 6 List of immune genes abbreviations**

| **Immune gene** | **Abbreviation** |
| --- | --- |
| Activated B cells | B Cells+ |
| Activated CD4 T cells | CD4 T cells+ |
| Activated CD8 T cells | CD8 T cells+ |
| CD56bright natural killer cells | CD56bright NK cells |
| CD56dim natural killer cells | CD56dim NK cells |
| Central memory CD4 T cells | C-Memory CD4 T cells |
| Central memory CD8 T cells | C-Memory CD8 T cells |
| Effector memory CD8 T cells | E-Memory CD8 T cells |
| Effector memory CD4 T cells | E-Memory CD4 T cells |
| Eosinophils | Eos |
| Gamma delta T cells | γδ T cells |
| Immature B cells | Immature B cells |
| Immature dendritic cells | Immature DCs |
| Myeloid-derived suppressor cells | MDSCs |
| Macrophages | MΦs |
| Mast cells | MCs |
| Monocytes | Monos |
| Memory B cells | Memory B cells |
| Natural killer T cells | NK T cells |
| Natural killer cells | NK cells |
| Neutrophils | PMNs |
| Plasmacytoid dendritic cells | PC DCs |
| Regulatory T cells | Tregs |
| T follicular helper cells | Tfh cells |
| Type 1 T helper cells | T1-Th cells |
| Type 17 T helper cells | T17-Th cells |
| Type 2 T helper cells | T2-Th cells |

**Supplementary Table 7 List of immune cells abbreviations**

| **Immune cell** | **Abbreviation** |
| --- | --- |
| Activated Natural killer cells | NK cells+ |
| Activated dendritic cells | DCs+ |
| Activated mast cells | MCs+ |
| Activated memory CD4 T cells | CD4 memory T cells+ |
| Eosinophil | Eos |
| Gamma delta T cells | γδ T cells |
| Macrophage M0 | M0-MΦs |
| Macrophage M1 | M1-MΦs |
| Macrophage M2 | M2-MΦs |
| Monocyte | Monos |
| Naive B cells | Naïve B cells |
| Naive CD4 T cells | CD4 naïve T cells |
| Neutrophil | PMNs |
| Plasma cells | PCs |
| Regulatory T cells | Tregs |
| Resting Dendritic cells | DCs- |
| Resting Natural killer cells | NK cells- |
| Resting mast cells | MCs- |
| Resting memory CD4 T cells | CD4 memory T cells- |
| T follicular helper cells | Tfh cells |

**Supplementary Figure 1 Boxplots comparison of leukocytes metagenes and cell fractions for predicted risk groups. Samples who couldn't be predicted were removed. (A)** Leukocytes gene expression **(B)** Leukocyte cell fractions. P-Values were obtained from Student's t-test. P-Value significance: * < 0.05, ** < 0.01.

**Fig 1A**

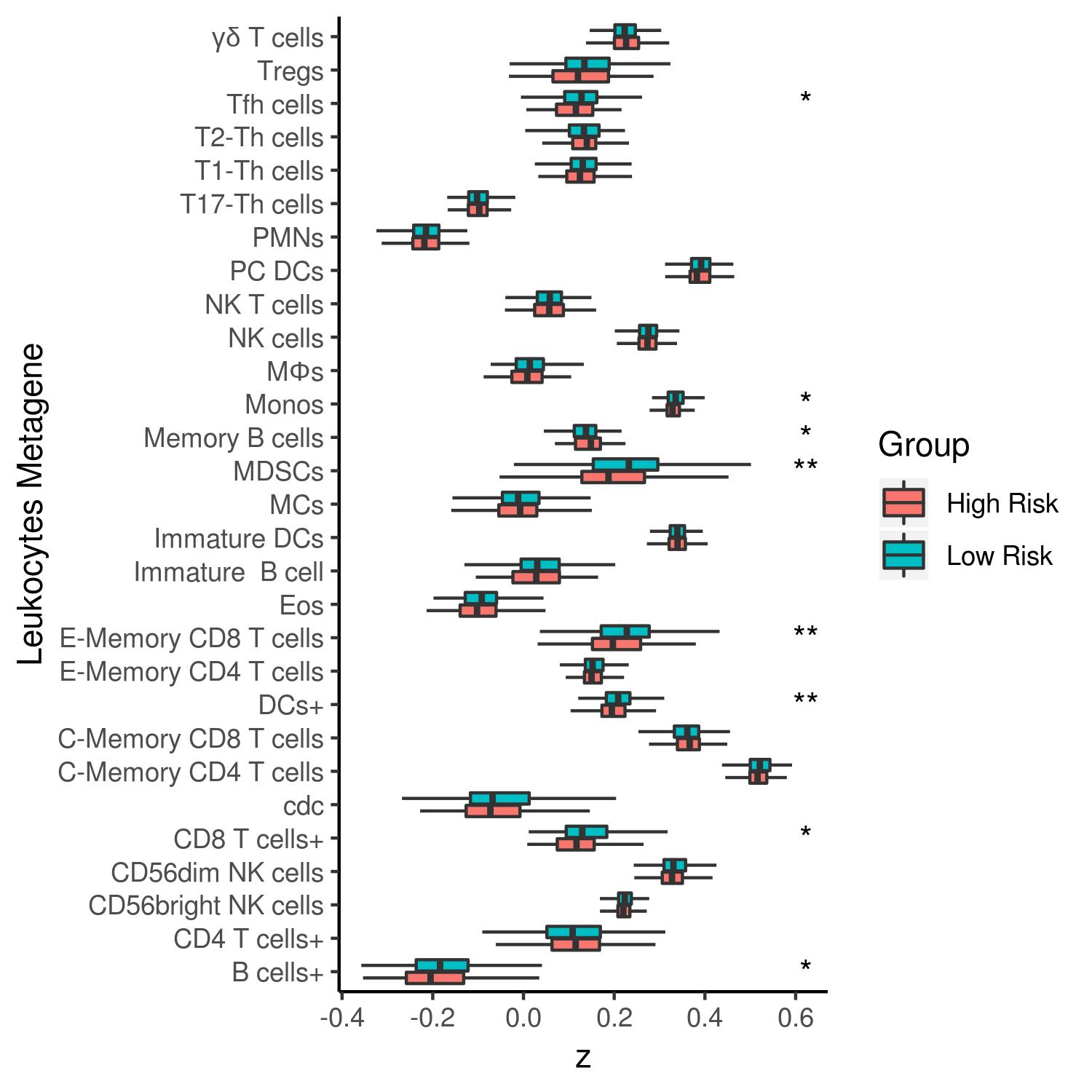

**Fig 1B**

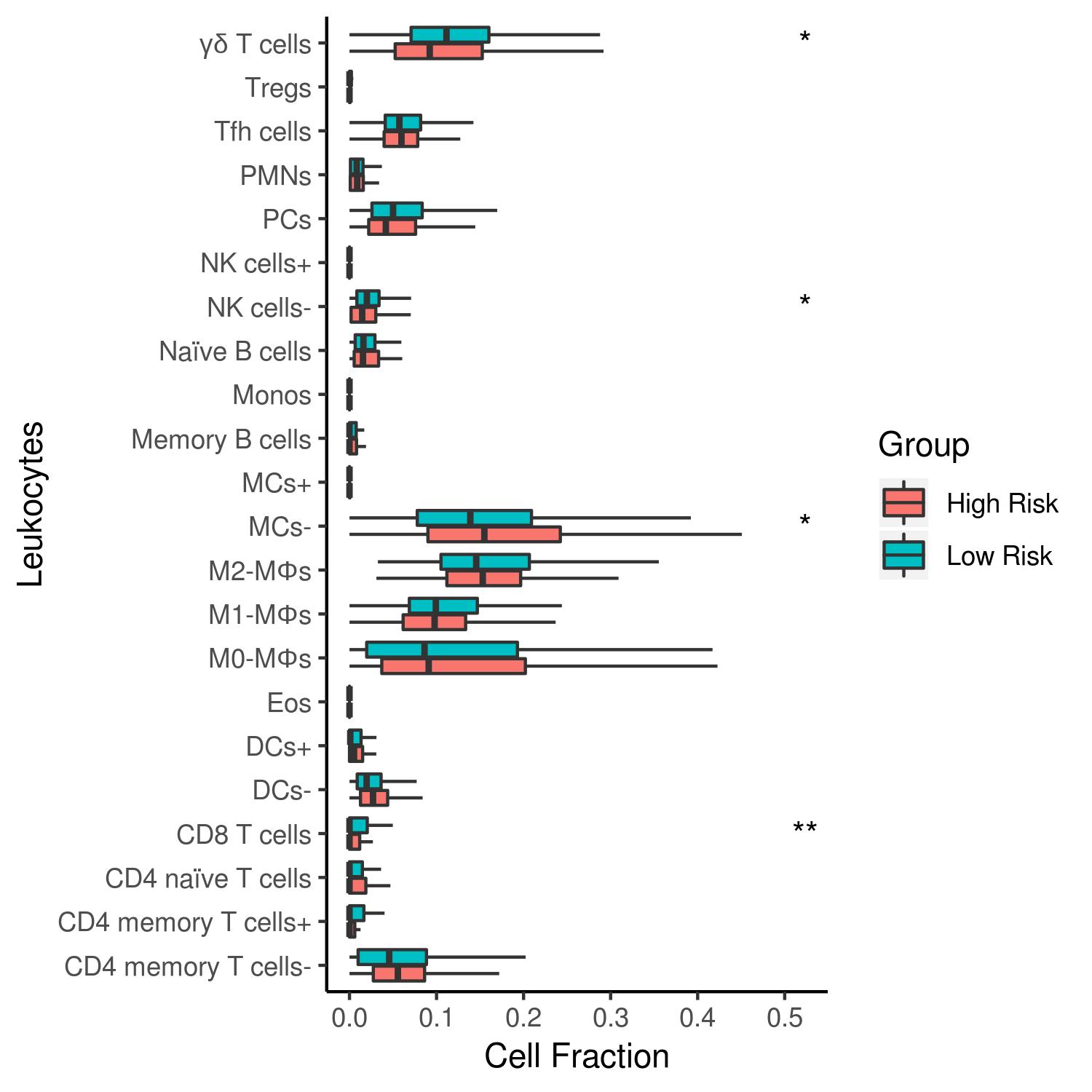

**Supplementary Methods**

**Sequencing data pre-processing and variant calling**

The GATK^1^ pipeline-based whole exome-sequence data pre-processing was described previously^2^. Briefly, duplicate reads were marked and removed using GATK’s markDuplicates removed using BamTools. Reads with a low mapping quality (n=60) were also removed using BamTools^3^. Local realignment around indels was made using GATK’s IndelRealign/RealignTargetCreator and finally, base recalibration was conducted using GATK’s BaseRecalibrator. All the variants were obtained using the Varscan2^4^ somatic option by analyzing normal/tumor matched sequencing files. Variants with strand-specific bias, coverage less than 30 reads and variant frequency for heterozygous calls of less than 0.08 were removed.

**Determine tumor purity**

Tumor purity was obtained using absCNseq^5^. To run absCNseq, for a given tumor, we generated a segmentation file and a SNV (Single Nucleotide Variants) file. The segmentation file was generated by running Varscan2 using the standard protocol. Briefly, we ran VarScan2’s copyNumber on normal and tumor BAM files, and VarScan2’s copyCaller to adjust for GC content and finally applied circular binary segmentation. The SNV file was then transformed from the VCF file by running VarScan2. For some samples, absCNseq could give a few purity solutions. In this situation, the consensus purity was selected. The samples with purity greater than 70% were retained for downstream analyses. Ultimately, 755 ER+ breast tumor samples were available for further analysis.

**Training and validation sets for ER+ breast cancer dataset**

To identify gene signatures of ER+ breast cancer, we randomly selected 200 samples, which have follow-up time, as the training set (30 and 170 for recurred and non-recurred samples, respectively). By default, ~15% of ER+ breast tumors get recurred within 10 years^6^. Clinically, at present almost all of the ER+/luminal breast cancer patients receive tamoxifen treatment. However, tamoxifen treatment for the ‘real’ low-risk patients does not affect patients’ survival. To develop gene signatures for prognosis, we controlled the training set such that we tried to make sure that the selected ‘low-risk’ patients are ‘real low-risk’ patients by applying these rules: (1) the low-risk patients who have relatively longer survival in the cohort, (2) we further confirmed them by predicting them using the gene-expression-based prognostic signatures we developed previously^7^. This signature was developed using a cohort where the patients have not been treated with any chemotherapy. The predicting accuracy for low-risk ER+ breast cancer reached 95%. Except these training samples, the rest of the ER+ breast tumor samples in the GDC was used for validation. Sixty non-recurred samples were retained for obtaining optimal signature cutoffs (Supplementary Table 5). Finally, all the remaining ER+ samples were separated into 2 validation sets (TCGA-Nature, TCGA-CPTAC) composed of 200 and 295 samples, respectively. For TCGA-Nature set, we used a ratio of 10% of recurred samples (20 recurred and 180 non-recurred samples). For TCGA-CPTAC, we used a ratio of 11.5% of recurred samples (34 recurred and 261 non-recurred samples).

**Leukocytes mutations comparative analysis**

To further assess leukocytes variants predictive power, we re-ran eTumorMetastasis^17^ pipeline by using only germline functional variants located in leukocytes metagenes^18^. As a comparative analysis, we used the same training and validation sets as well as the same cutoffs used previously with all functional germline variants. Gene sets were generated randomly for each GO Terms (Apoptosis, Cell Cycle, Cell Adhesion, Cell Proliferation, Immune Response and Cell Proliferation) and then we applied a multiple survival screening algorithm^7,17^. For each GO Term, not enough significant gene sets were obtained for us to be able to extract a gene signature.
